# Systematic identification of seed-driven off-target effects in Perturb-seq experiments

**DOI:** 10.64898/2026.03.27.714658

**Authors:** Austin Hartman, John D. Blair, Thao P. Nguyen, Kyle Dyson, Alexandra Bradu, Oliver Takacsi-Nagy, Katherine Santostefano, Theresa Boade, Matthew Bolaños, Ronghui Zhu, Emma Dann, Alexander Marson, Aaron D. Gitler, Rahul Satija, Ansuman T. Satpathy, Theodore L. Roth

## Abstract

Genome-wide Perturb-seq (GWPS) has emerged as a powerful approach for unbiased mapping of gene regulatory networks. A key assumption underlying many Perturb-seq analyses is that each guide RNA exclusively perturbs a single target locus. Without methods to identify and filter off-target events, erroneous gene-pathway associations driven by off-target activity can propagate into downstream analyses. Here, we present a workflow for the systematic identification of candidate off-target events in CRISPRi Perturb-seq experiments. Our approach exploits the observation that cells harboring a guide which represses an off-target gene display transcriptional similarity to cells in which that gene is directly targeted by an on-target guide. We apply our workflow to multiple GWPS datasets and nominate off-target events in which a guide nominally targeting one gene also represses a distinct gene producing a phenotype likely attributable to the off-target perturbation. We use both off-target gene repression and guide seed sequence alignments at the off-target promoter locus as evidence for off-target effects and find independent evidence of putative off-target events in separate GWPS datasets. Together, these results establish a principled framework for the identification and filtering of off-target guide effects in Perturb-seq experiments.

## Main

GWPS screens, which combine pooled CRISPR perturbations with single-cell RNA-seq, have emerged as a powerful approach for comprehensive mapping of gene regulatory networks^1–3^. A foundational assumption underlying most Perturb-seq analyses and methods is that each guide mediates perturbation exclusively at a single target locus. Individual cases of guides producing off-target effects are well-studied^4–8^, but there are few options for systematic identification of off-target guides and the resulting transcriptional effects in Perturb-seq experiments. Commonly used genome-wide guide libraries, including Dolcetto, Brunello, Calabrese, and GeCKO were designed to maximize on-target activity while minimizing off-target activity^9–15^. The recently designed Katsano CRISPRi Cas9 guide library specifically aims to minimize off-target effects derived from seed matches^16^. In addition to guide design, researchers have explored additional CRISPR systems^17,18^ and variants of Cas9^19–22^, in part, to further optimize the sensitivity and specificity of CRISPR targeting systems. Although off-target activity is theoretically minimized at the guide library design stage, experimental validation at large scale remains difficult, particularly because off-target effects may be cell context-dependent. Consequently, guides with off-target activity likely remain present in genome-wide guide libraries.

Recent reports indicate the importance of the seed-region (the bases most-proximal to the protospacer adjacent motif or PAM) of guide RNAs in dCas9 recruitment in CRISPRa screens^23,24^. Thus, we sought to determine whether sequence complementarity within the PAM-proximal seed region of guides could account for, and ultimately predict, off-target transcriptional repression in CRISPRi screens. Because short PAM-adjacent DNA sequences occur with high frequency throughout the genome and promoter loci make up a small fraction of the genome, an exhaustive search of all potential seed matches would yield a large number of candidate sites–many of which are unlikely to cause a substantial biological effect. We therefore reasoned that leveraging transcriptional readouts from existing Perturb-seq datasets would provide a means of filtering the search space, restricting our analysis to off-target events with direct functional evidence and relevance for the interpretation of genome-wide screens.

In this study, we developed a systematic approach to identify candidate off-target events in GWPS datasets. We applied this approach to datasets utilizing different guide libraries across multiple cell types. We observed a trend between the length of a seed match at an off-target locus and the magnitude of observed transcriptional repression. We next examined characteristics of off-target guides, including sequence composition and position relative to the off-target TSS. Applying these rules to additional Perturb-seq screens, we identified likely off-target effects among genes implicated in TCR signaling, which were restricted to guides with off-target seed matches near the TSSs of canonical TCR signaling genes and absent for independent guides targeting the same genes.

## Results

We designed a workflow that consisted of three steps: (1) guide clustering, (2) guide seed alignment at candidate genes, and (3) filtering for transcriptional repression at candidate genes (**Fig. 1a**). Co-clustering of perturbations in a common pathway has been documented across multiple GWPS experiments and methods^25–30^. Thus, the first step of our pipeline rests on the assumption that guides targeting members of a common pathway tend to result in similar transcriptomic changes when knocked down and are therefore nearby in low-dimensional embeddings of guide pseudobulk profiles. Critically, we reasoned that guides which incidentally repress a member of a given pathway through off-target activity would also cluster with on-target guides for that pathway. Building on this logic, we constructed guide neighborhoods by first aggregating single-cell profiles at the guide level via pseudobulking and subtracting the non-targeting guide mean from each pseudobulk vector. Dimensionality reduction was performed using PCA with 100 principal components (**Methods**). For each guide, the 20 nearest neighbors in the embedding were identified. We then extended each neighborhood to include guides not among the 20 immediate neighbors, but which themselves list the guide among their own nearest neighbors. Asymmetrical neighbors were included to account for asymmetric off-target effects: a guide may cluster primarily with others targeting the same pathway yet exert enough off-target repression of a gene in a second pathway to also appear among that pathway’s guides. Second, we searched for candidate off-target binding sites near TSSs of genes targeted by neighboring guides. This search aligned guide seed regions, defined as the 10-12 PAM-proximal bases^16^, using a more permissive threshold of ≥5 contiguous matching bp adjacent to a PAM to candidate TSSs. Finally, for each neighboring guide in a guide neighborhood we asked whether it repressed any of the genes targeted by the neighboring guides. While in many cases this may be biological, we hypothesized that a subset of cases occurred due to off-target repression of the gene targeted by the neighboring guide, resulting in neighborhood membership. Ultimately, we generated a list of guides with both transcriptomic evidence of off-target repression, and a candidate locus mediating the effect.

**Figure 1:**
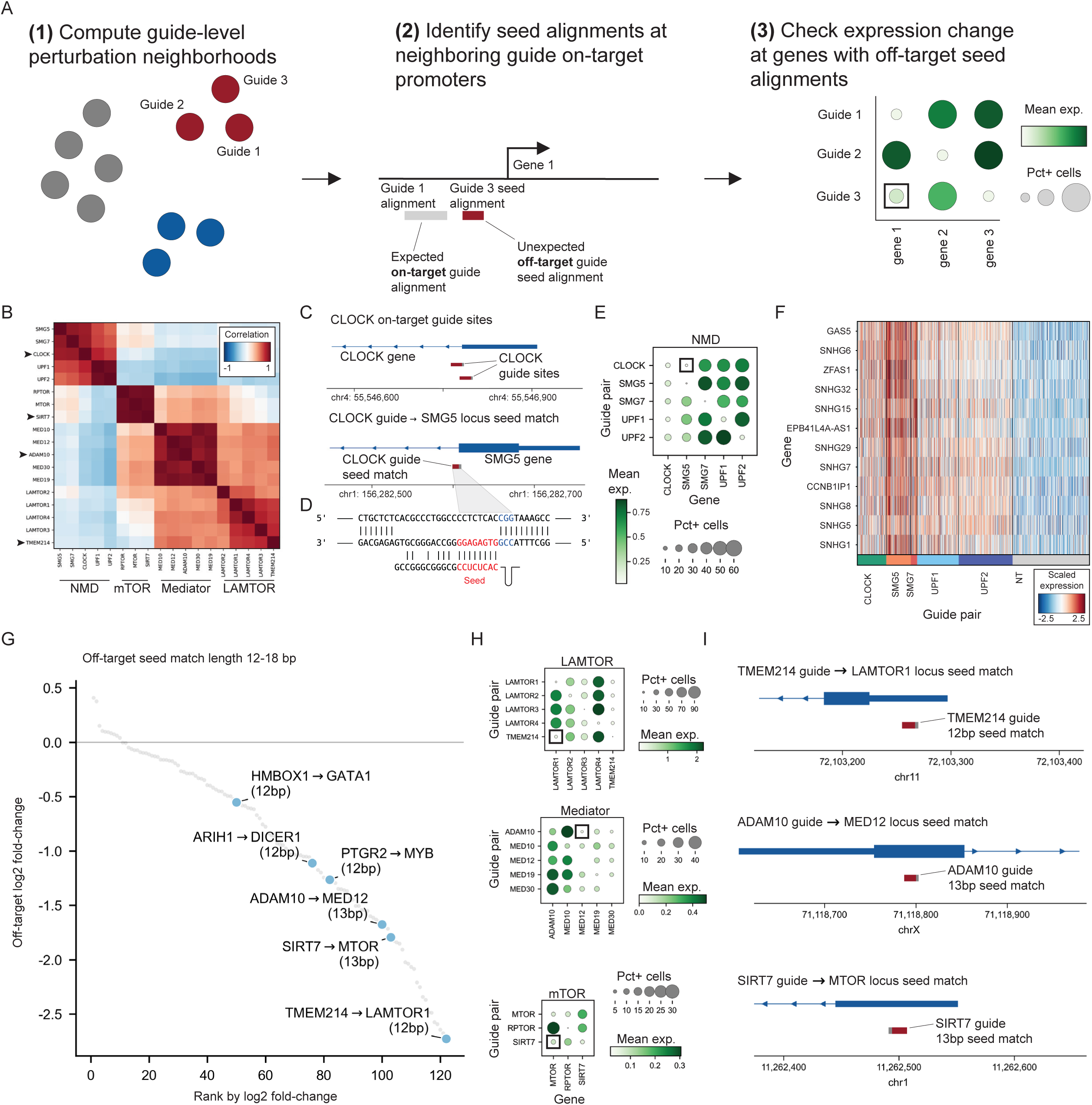
Identification of candidate off-target guides. **a.** High-level overview of the workflow for identification of off-target guide candidates. **b.** Heatmap displaying pseudobulked correlation for guides designed to target genes within the mediator, NMD, mTOR, and LAMTOR pathways or candidate off-target guides with transcriptional similarity (See methods). **c.** Loci surrounding the *SMG5* and *CLOCK* TSSs. The *CLOCK* locus is annotated with the alignments of each *CLOCK* guide. The *SMG5* locus is annotated with a candidate off-target binding site in which the first 8 bp of *CLOCK* guide 1 align directly adjacent to a PAM site. **d.** Alignment of candidate off-target *SMG5* locus and *CLOCK* guide #1. Bars indicate complementarity between the locus and guide sequence. **e.** Dot plots displaying normalized expression of the *CLOCK, SMG5, SMG7, UPF1,* and *UPF2* genes across cells receiving guides designed to target *CLOCK, SMG5, SMG7, UPF1,* and *UPF2*. **f.** Single-cell heatmap displaying the top 12 *CLOCK* DEGs relative to non-targeting controls by a Wilcoxon rank-sum test with a minimum log2 fold-change of 1 and a maximum p-value of 0.01 across *CLOCK*, non-targeting cells, and canonical members of the nonsense mediated decay pathway; *SMG5*, *SMG7*, *UPF1*, and *UPF2*. NT is a random sample of 500 cells receiving non-targeting control guides. **g.** Log2 fold-change rank plot of identified seed matches between 12-18 bp. Labels indicate the guide target with an arrow pointing to the predicted off-target gene. The numbers in parenthesis indicate the number of matching bp between the off-target locus and seed, not including the PAM. **h.** Dot plots displaying the expression of canonical pathway members and candidate off target guides. **i.** Loci surrounding the *MED12*, *LAMTOR1*, and *MTOR* TSSs annotated with candidate off-target guide seed alignments (in red), adjacent to PAM sequences (in grey). Seed matches were 12, 13, and 13 bp for *TMEM214, ADAM10*, and *SIRT7* guides, respectively.

To validate our pipeline, we applied it to a GWPS dataset in K562 cells and asked whether our pipeline identified previously proposed off-target guide candidates described in the original work^25^. The first case involved a guide pair targeting *CLOCK*; cells receiving *CLOCK*-targeting guides shared transcriptional similarity with cells receiving guides targeting canonical nonsense-mediated decay (NMD) components, including *SMG5*, *SMG7*, *UPF1*, and *UPF2*^25^. In a pseudobulked PCA embedding (**Methods**), *CLOCK*-targeting guides were grouped with guides targeting canonical NMD factors *SMG5*, *SMG7*, *UPF1*, *UPF2* (**Fig. 1b**)^31^. We then searched for seed matches near the neighboring genes, reasoning that this would establish a plausible locus for off-target recruitment of dCas9. An 8 bp seed match adjacent to an NGG PAM was identified for one of the *CLOCK* guides near the *SMG5* TSS with 6 additional matches in the remaining 12 bp of the guide (**Fig. 1c,d**). Consistent with prior observations, cells that received the *CLOCK* guide exhibited significant downregulation of both the intended target, *CLOCK,* and the NMD factor, *SMG5,* but not other members of the NMD pathway (**Fig. 1e**). Further, we found that differentially expressed genes in cells receiving the *CLOCK* guide pair were also differentially expressed after knockdown of canonical members of the NMD pathway (**Fig. 1f**). We applied our pipeline to the entire K562 GWPS dataset and nominated over 100 additional candidate off-target events^25^, each supported by repression of a neighboring gene and the presence of a seed match of ≥12 bp adjacent to a PAM (**Fig. 1g**). Our analysis revealed additional examples, such as a guide pair targeting *TMEM214*, whose recipient cells exhibited repression of *LAMTOR1,* as previously reported in the K562 GWPS paper^25^. We recapitulated this observation and identified a 12 bp match at a PAM-proximal site near the *LAMTOR1* TSS (**Fig. 1h,i**). Among additional putative off-target effects identified, we highlighted possible off-target repression of *MED12* by an *ADAM10* guide, and putative off-target repression of *MTOR* by a *SIRT7* guide, each supported by 13 bp seed matches at the off-target loci and downregulation of the respective off-target gene (**Fig. 1h,i**). Collectively, these results demonstrate the utility of this approach for systematic identification of candidate off-target activity, with particular emphasis on perturbations that may confound the interpretation of downstream functional analyses.

### Application to additional genome-wide Perturb-seq datasets

To assess the generalizability of our pipeline, we applied it to three GWPS datasets spanning distinct guide libraries, cell lines, and cell types: (1) K562 cells [Replogle] with dual-targeting guides (two guides targeting the same gene on the same vector), (2) HCT116 cells [Xaira] with dual-targeting guides, and (3) CD4+ T cells [Zhu & Dann] with two independent guides per gene (**Extended Data Fig. 1a**)^25–27^. All three datasets utilized CRISPRi methods that tether dCas9 to a ZIM3 Krüppel-associated box (KRAB) domain, a C2H2 zinc-finger protein domain which emerged as a potent transcriptional repressor in a screen of diverse KRAB domains (**Extended Data Fig. 1b**)^32^. Across these datasets, we identified thousands of cases with seed region complementarity of ≥5 bp within the promoter of a co-repressed neighboring gene. We selected 5 as the minimum seed length match, as previous reports have identified dCas9 recruitment to loci with as few as 5 bp seed matches^23,24^. Strikingly, seed match length was positively associated with the magnitude of transcriptional repression, with effects becoming apparent at seed matches of ∼10 bp (**Fig. 2a**). We attributed this trend to two complementary factors: first, shorter seed matches are expected to harbor a substantially higher false-positive rate, as short sequence motifs occur at greater frequency throughout the genome, and second, we hypothesized that longer seed matches mediate stronger recruitment of dCas9 to the off-target locus, thereby mediating stronger repression. Accordingly, seed matches of 12 bp or more frequently repressed targets at levels comparable to on-target guides (**Fig. 2a,b**)^33^.

**Figure 2:**
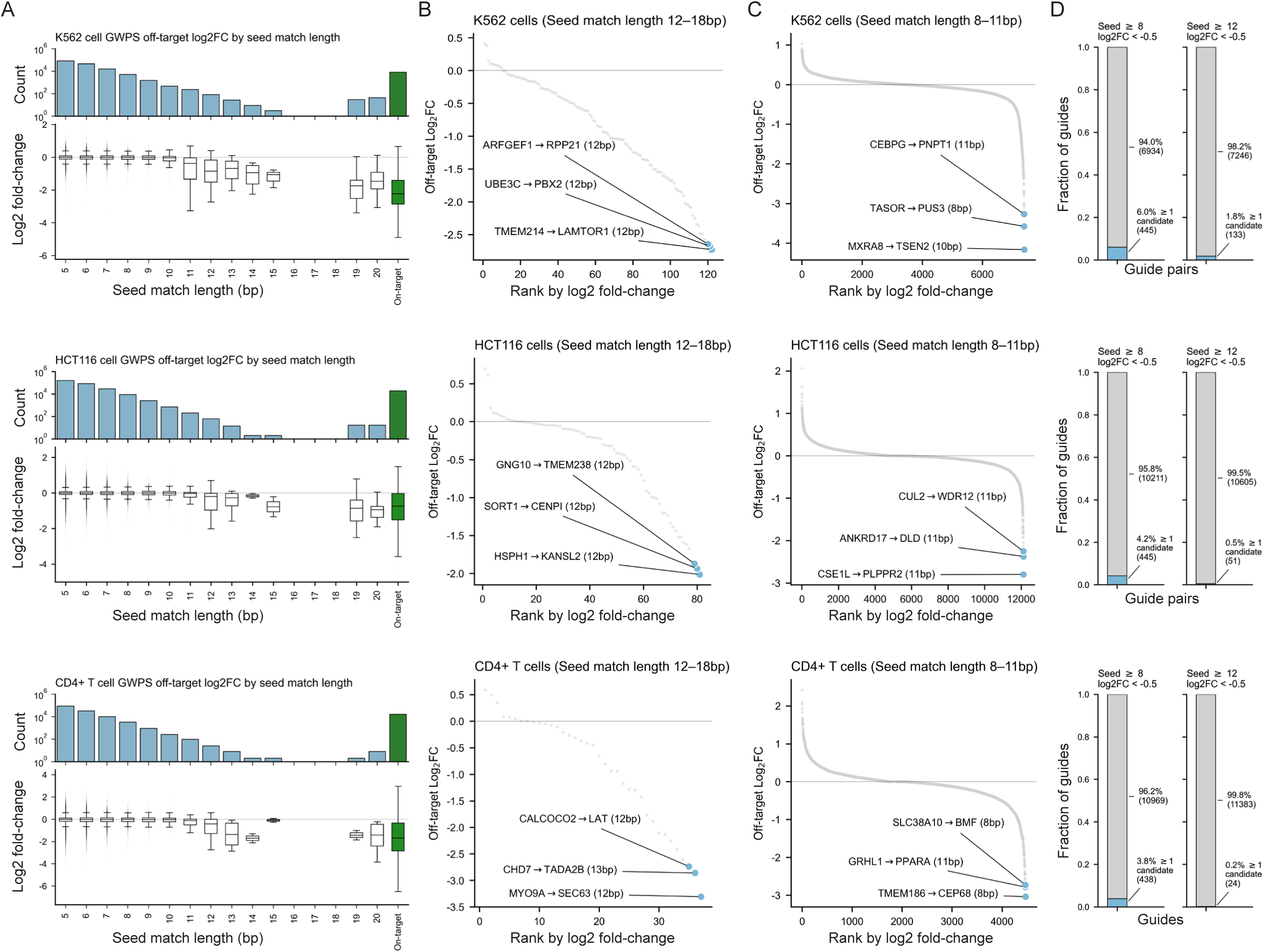
Nomination of candidate off-target guides across genome-wide Perturb-seq data. **a.** Measured log2 fold-change for candidate off-target guides at the predicted off-target guide split by the length of seed sequence match. Seed matches of 19 or 20 bp indicate off-target repression of a neighboring gene, while the others represent predicted distal off-target events. The final column of the plots displayed measured log2 fold-change for guides at their intended targets. **b.** Log2 fold-change rank plot showing off-target 12-18 bp seed matches across three GWPS datasets. **c.** Log2 fold-change rank plot showing off-target 8-11 bp seed matches across three GWPS datasets. **d.** Bar plots showing the fraction of affected guides, of guides with on-target log2 fold-change knockdown greater than 0.5 and greater than 20 cells, which the workflow uses as a cutoff. The first bar plot shows the fraction of those guides with an 8 or more bp seed match within 2 kb of an off-target gene with greater than 0.5 repression. The second bar plot shows the same for seed matches of 12 bp or more.

Notably, all candidate off-target events with 19-20 bp seed matches represented cases of off-target repression of genes whose TSSs lie in close proximity to the on-target gene TSS. We attributed the relative scarcity of candidates with ≥14 bp matches to design constraints of guide libraries, which enforce a minimum Hamming distance between guide sequences and potential off-target sites across the genome. To isolate high confidence candidate off-target events, we subsetted to seed matches of 12 bp or more (and excluded matches of 19 and 20 bp representing repression of a nearby gene) and observed that a majority of cases resulted in lower expression of the candidate off-target gene (**Fig. 2b**). Additionally, given that shorter seed matches may lead to off-target activity, we examined the log2 fold-change for seed matches between 8 and 11 bp which also showed evidence of transcriptional repression, but to a lesser degree than seed matches of 12-18 bp (**Fig. 2c**). Lastly, we computed the fraction of guides in each guide library which are nominated for off-target activity. 4.67% (range: 3.8-6.0%) of guides contained a seed match greater than or equal to 8 bp and downregulation of the off-target gene. 0.83% (range: 0.2%-1.8%) of guides were affected when we filtered to only include seed matches of 12-18 bp (**Fig. 2d**). Although there were differences in the number of candidate off-target events in each dataset, we note that the number of off-target events identified in a dataset should not be interpreted as a measure of data quality, as it reflects several factors including the number of guides, guide design, number of cells per guide, expression levels of targeted genes, and active pathways within cells (**Extended Data Fig. 1f**).

We additionally applied our workflow to a VIPerturb-seq dataset^29^ (**Extended Data Fig. 1c**), which uses a CRISPRi dCas9-KRAB-MeCP2 cassette^34^ that is different from each of the other three GWPS datasets (**Extended Data Fig. 1b**). We detected fewer off-target candidates, particularly with ≥12 bp seed matches, but it is likely that lower cell recovery per guide limits the sensitivity of detection (**Extended Data Fig. 1c,d**). For the top three candidates, off-target repression was detected by the predicted guide, and not by two additional guides targeting the same on-target gene (**Extended Data Fig. 1e**). For the CD4+ T cell dataset, we generated our predictions by analyzing one of twelve publicly available GWPS datasets spanning 4 donors and 3 conditions. We further validated these predictions generated using donor 1 (**Fig. 2d**) using data from the 3 additional donors and observed a consistent pattern of off-target repression (**Extended Data Fig. 2b**). To further assess the reproducibility of candidate off-target events, we validated guide pairs identified in the K562 cell^25^ experiment using a separate GWPS experiment in HCT116 cells^27^ with many of the same guide pairs. In the HCT116 dataset, we similarly observe candidate off-target repression of the *LAMTOR1*, *MED12*, and *MTOR* genes by the same guides (**Extended Data Fig. 2c**).

### Characteristics of candidate off-target guides

To further understand why certain guides appeared to have off-target activity, we explored the sequence characteristics of nominated off-target guides. To do so, we selected candidate off-target guides with ≥10 bp off-target seed matches and computed the positional base frequency relative to the entire guide library. Across datasets and guide libraries, we observed an enrichment in guanines in the 10 bases proximal to the PAM (**Fig. 3a**) as well as an enrichment in PAM motifs, relative to other guides in the library as others have reported (**Methods**, **Extended Data Fig. 3a**)^16,23^.

**Figure 3:**
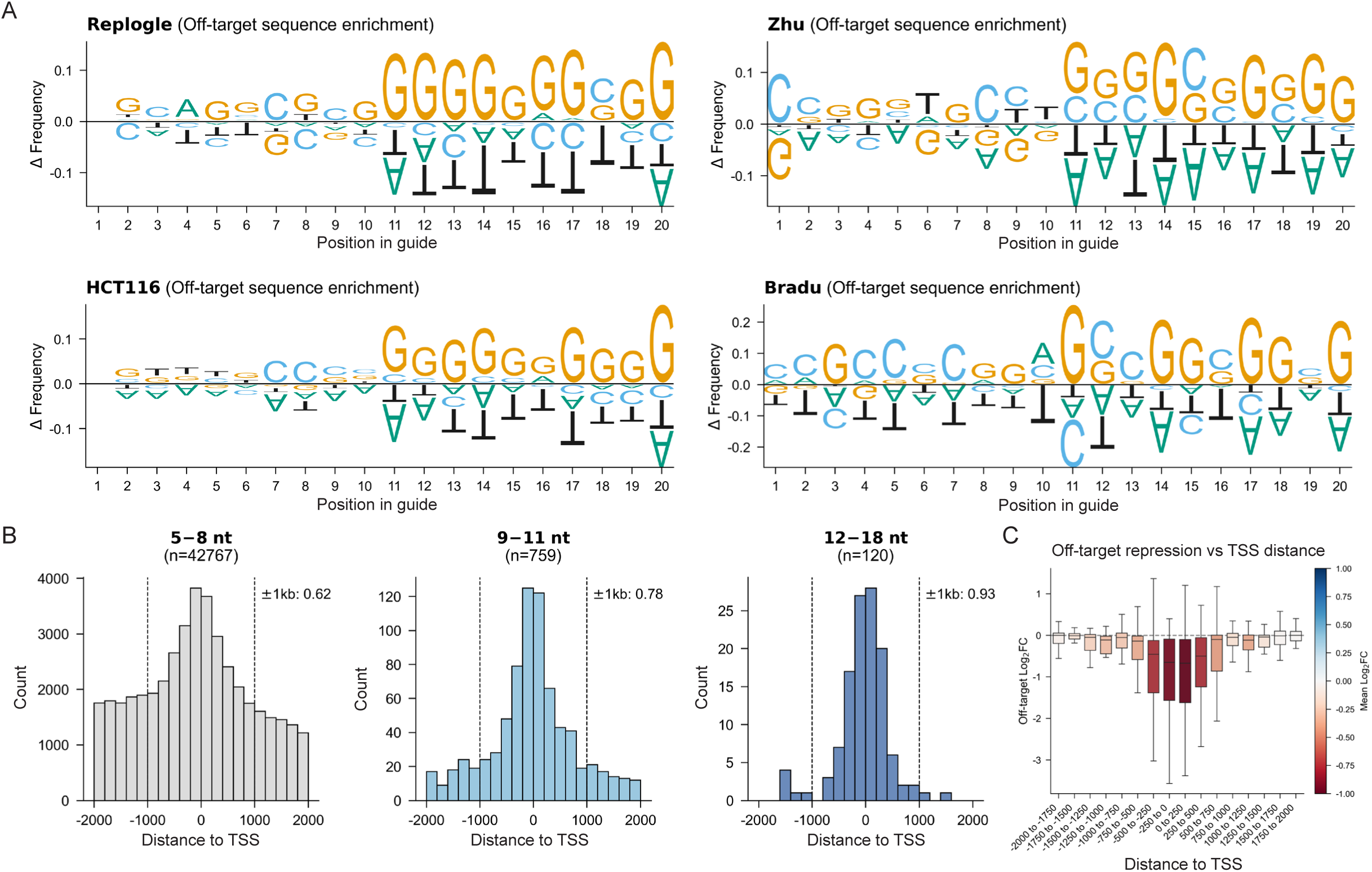
Sequence characteristics of candidate off-target guides. **a.** Plots displaying the difference in base-frequency at each guide position for candidate off-target genes, relative to the full guide libraries. Candidates were defined as seed matches of 10-18 bp with a negative log2 fold-change across four GWPS datasets. **b.** Histograms of off-target seed alignment position relative to off-target TSS grouped by length of seed match across four GWPS datasets. The number of candidate events involving a guide and off-target locus were shown in parenthesis and the fraction of those events within +/-1,000 bp of the annotated TSS were labeled. **c.** Magnitude of log2 fold-change repression by seed alignment position relative to the off-target TSS. Candidate sites include all seed matches of 11-18 bp, without filtering on log2 fold-change.

To investigate the relationship between off-target guide RNA activity and genomic context, we visualized candidate off-target sites relative to the off-target TSS across four GWPS datasets^25–27,29^. We hypothesized that seed region complementarity exceeding 12 bp would confer a higher probability of functional dCas9 recruitment compared to off-target loci with shorter seed matches of 5 to 8 bp. Consistent with this, TSS enrichment was markedly stronger among high-confidence candidates (seed match ≥12 bp and log2 fold-change < -0.5) (**Fig. 3b**). Notably, even among candidates with shorter seed matches, where false-positive rates are expected to be elevated, we still observed significant enrichment within approximately 1,000 bp of the annotated TSS. However, this may reflect the elevated GC content characteristic of promoter sequences. Finally, to assess the functional consequences of off-target recruitment, we quantified the magnitude of transcriptional repression at candidate off-target genes as a function of signed guide-TSS distance. This revealed a moderate but consistent inverse relationship, supporting a model in which the magnitude of dCas9-KRAB-mediated repression is partly determined by proximity to the TSS (**Fig. 3c**)^35^. Our analysis revealed that long seed matches near off-target TSSs are a likely source of off-target CRISPRi activity. Given this, we computed all possible seed matches of 6-15 bp within 1,000 bp of protein coding TSSs and made this available as a web application to help evaluate the specificity of new and existing CRISPRi guides (**See code & resource availability**). We then quantified the fraction of guides in genome-wide CRISPRi libraries with off-target seed matches ≥12 bp within 1,000 bp of a protein-coding TSS and found that 47.2–59.8% of guides had at least one match (**Extended Data Fig. 3b**). However, we emphasize that not all seed match events will result in an effect due to additional factors such as chromatin state and expression of the off-target gene.

### Candidate off-target events may lead to erroneous association with TCR signaling

Finally, we asked whether seed matches may complicate the interpretation of additional GWPS experiments. A recent preprint described GWPS experiments in Jurkat cells and nominated several novel candidate regulators of the TCR pathway^36^. The authors trained a perturbation prediction model and fine-tuned it using GWPS data from resting Jurkat cells. The authors then asked whether it could accurately predict perturbation responses in activated Jurkat cells and suggested that their predictions were closer to ground truth than a linear baseline. We analyzed the guide library for off-target seed matches and found that three of the seven nominated candidate regulators–*LRBA, APPL2* and *WDR53*–had off-target seed matches at the promoters of *LAT* and *CD3D*, two known TCR pathway genes, possibly resulting in misassociation with TCR signaling (**Fig. 4a**). We also noted that the *APPL2* and *WDR53* guides had 83% GC-content in the 12 bp seed region, and the *LRBA* guide contained 5 consecutive guanines (**Fig. 4a).** To investigate whether the impact on TCR signaling was due to on-target repression of *LRBA*, *APPL2* and *WDR53* or off-target repression of *LAT* and *CD3D*, and since this dataset was not available for re-analysis, we examined the prior primary CD4+ T cell dataset^26^. The CD4+ T cell guide library included two guides for *APPL2*–one of which was the same guide as in the Jurkat GWPS dataset with *LAT* promoter homology–and two independent guides targeting *WDR53*. The *LRBA* guide with multiple *CD3D* seed alignments was also shared between the CD4+ T cell and Jurkat GWPS guide libraries. We found that only the shared *APPL2* guide reduced *LAT* levels, while both guides effectively repressed *APPL2* (**Fig. 4b,c**). Further, neither of the independent *WDR53* guides displayed the TCR inactivation phenotype, which was marked by increased levels of *IL32, MAL, GZMA*, and *LTB* in the primary CD4+ T cell dataset (**Fig. 4b,c**). Importantly, the TCR inactivation phenotype was observed exclusively for the *APPL2* and *LRBA* guides with *LAT* and *CD3D* seed matches and not for the remaining four guides (*WDR53*-1, *WDR53*-2, *LRBA*-1, and *APPL2*-1), despite repression of their intended targets (**Fig. 4b,c**).

**Figure 4:**
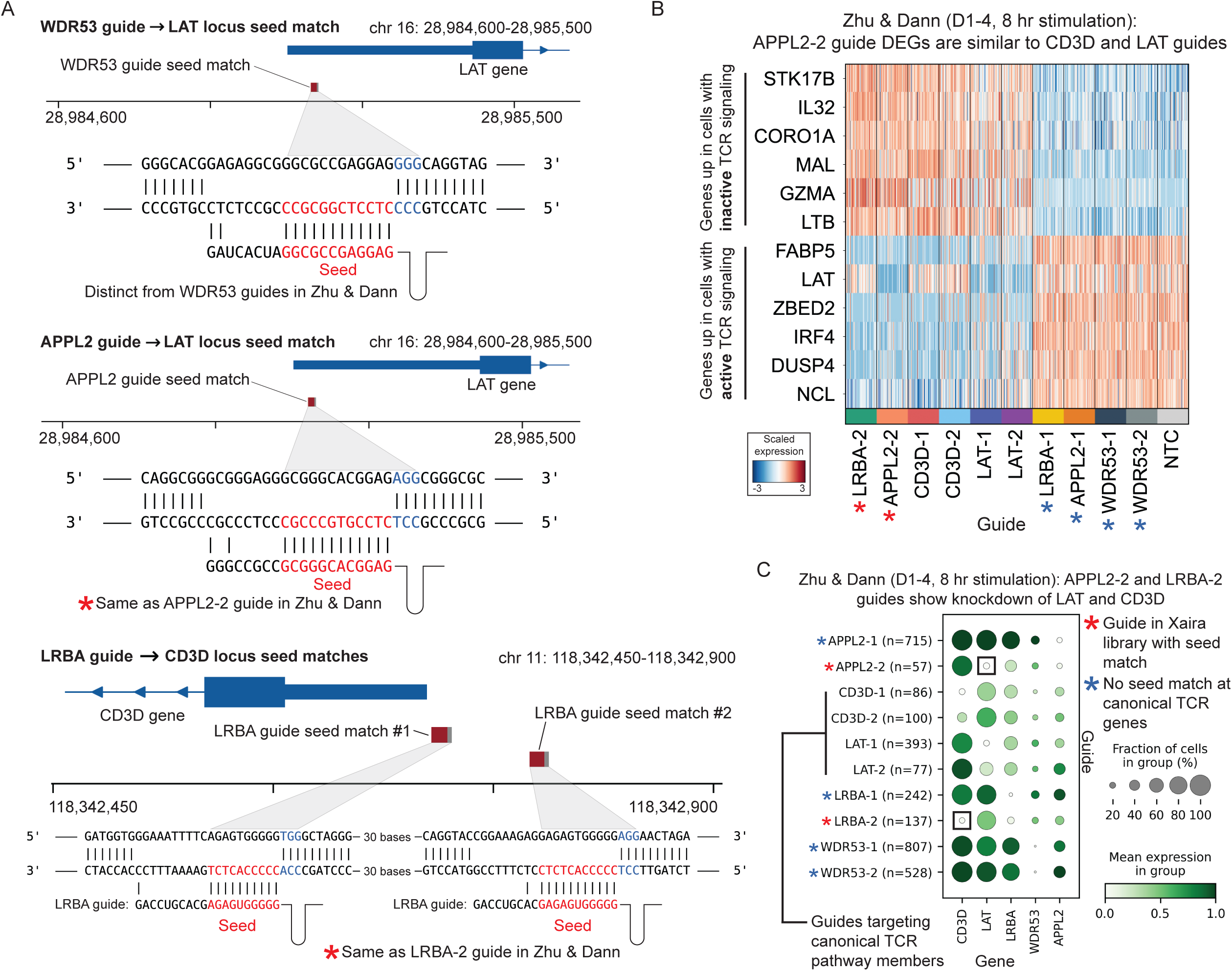
Off-target seed matches may drive false associations with TCR signaling. **a.** Loci surrounding the *LAT* and *CD3D* TSSs and PAM-adjacent seed matches surrounding the loci for *APPL2*, *WDR53,* and *LRBA*. Alignments of the whole guide sequences are displayed below as well as the adjacent PAM sequences. **b.** Heatmap displaying top DEGs between cells receiving the APPL2-2 guide and non-targeting control guides. Blue asterisks indicate guides which do not contain seed alignments to *LAT* or *CD3D*. Red asterisks indicate guides found in the Jurkat GWPS and primary CD4+ T GWPS guide libraries with seed alignments to the *LAT* and *CD3D* loci. **c.** Dot plot of *CD3D*, *LAT*, *LRBA, WDR53*, and *APPL2* genes across cells receiving *APPL2*, *CD3D*, *LAT*, *LRBA,* and *WDR53* guides.

To further validate whether these phenotypes were guide-specific and likely due to off-target repression, we reanalyzed several independent genome-wide CRISPR screens in T cells^37–39^. In CRISPRi screens for IFNγ regulators in CD8+ T cells and IL-2 regulators in CD4+ T cells (two cytokines downstream of TCR activation), canonical TCR signaling members were consistently enriched as negative regulators, whereas *WDR53*, *LRBA*, and *APPL2* were not (**Extended Data Fig. 4a**). The candidate off-target guides for *WDR53*, *LRBA*, and *APPL2* were each absent from these libraries, likely explaining why these genes were not enriched as negative regulators. However, a *CALCOCO2* candidate off-target guide present in the CD4+ T cell GWPS library but not the Jurkat GWPS dataset was also represented in the CRISPRi experiments where it showed modest enrichment (**Extended Data Fig. 4a**). To investigate this further, we requantified enrichment at the guide level rather than at the gene level. Only the candidate off-target *CALCOCO2* guide showed enrichment, though to a lesser degree than the guides directly targeting the gene it is predicted to repress (**Extended Data Fig. 4b**). In summary, while canonical TCR pathway members were consistently enriched across CRISPRi, CRISPRa, and CRISPR knockout screens in both human and mouse T cells, *WDR53*, *LRBA*, and *APPL2* failed to show similar enrichment—suggesting that their appearance in the Jurkat GWPS dataset likely reflects off-target effects rather than genuine biological signal (**Extended Data Fig. 4c,d,e**).

## Discussion

Here, we describe a pipeline for filtering off-target effects in Perturb-seq datasets. The landscape of off-target events we identify varies substantially across datasets, which we suspect is not only a consequence of differences in guide library design, but also of cell-type-specific gene expression. For example, a guide with seed complementarity to the *LAT* promoter will likely fail to produce detectable knockdown or co-clustering with TCR pathway members in K562 cells where this pathway is transcriptionally silent yet may do so in T cells where the pathway is active. Analogous to recent work demonstrating that context-dependent perturbation responses can unmask novel biological regulators^40^, we anticipate that certain off-target events will only result in phenotypes in particular cellular contexts.

We believe that off-target effects will be an important consideration for machine learning models trained on large Perturb-seq atlases, where datasets for training often share a common guide library. In such cases, models may be systematically exposed to the same off-target confounders and learn spurious gene-pathway associations as genuine regulatory relationships. We therefore advocate for rigorous off-target filtering as a prerequisite for any computational framework that nominates pathway regulators from Perturb-seq data and emphasize the importance of reproducibility across multiple guides targeting the same gene. Further, recent Perturb-seq vector designs have utilized dual-guide approaches (two guides on the same vector) to enhance the likelihood and potency of target gene knockdown^25,27,36^. Future work should weigh these strategies alongside the increased risk of off-target effects, particularly for model training. Although the present analysis focuses on CRISPRi, we note that our pipeline may be flexibly applied to CRISPRa Perturb-seq experiments or CRISPRi experiments with different effectors with minor parameter adjustments.

While prior studies have investigated off-target activity of Cas9 nucleases^4–6,41–44^, dCas9 recruitment may require less guide homology to induce a transcriptional effect via the tethered effector. Even weak interactions at off-target loci could position the effector domain sufficiently close to a promoter or enhancer element to modulate transcription. This distinction is particularly relevant for CRISPRa and CRISPRi systems employing potent transcriptional activators or repressors, such as the VPR or KRAB domains^35,45,46^. The expression level of dCas9 may also influence the likelihood of repression at candidate off-target sites. However, we note that further experimental validation remains necessary, particularly for candidates supported by short seed matches and modest target gene repression, since dCas9 recruitment is likely influenced by genomic context including chromatin accessibility and epigenetic state which we do not take into account^16,47^. Thus, we imagine this approach could be further optimized with ATAC-seq and other measures of chromatin state^48^. Lastly, it may be useful to implement strategies for predicted non-seed-mediated off-target effects as recent work highlights that not all off-target effects appear to be driven by seed matches^16^. Overall, we hope this work will be utilized to mitigate the likelihood of spurious gene-pathway associations in large-scale GWPS datasets and to drive iterative improvement of genome-wide guide libraries that minimize off-target events.

## METHODS

### Guide neighborhood finding

Pseudobulk expression profiles were computed for each guide vector (Xaira and Replogle datasets deliver guides in pairs) by normalizing each cell to 10,000 counts, applying a log(x+1) transformation, and averaging across all cells assigned to each guide vector. The means for non-targeting guides are then subtracted from each targeting guide pseudobulk. The top 5,000 highest-variance features were selected for dimensionality reduction using PCA keeping the top 100 principal components. For each guide, the 20 nearest neighbors were identified using Euclidean distance. Additionally, neighborhoods for each guide are extended if a given guide appears in another guide neighborhood, but not its own.

### Guide seed search at candidate off-target promoters

For each potential off-target event, the promoter of the candidate off-target gene was extracted from the hg38 reference genome using a ±2,000 bp window (totaling 4,000 bp) surrounding the annotated TSS. Seed matches were identified as the longest contiguous match of ≥5 bp between the 3′ end of the spacer at any PAM-adjacent site within the window using string matching. For each candidate event, the seed match length, genomic coordinates, strand, and full-spacer Hamming distance between the guide sequence and the matched genomic site were recorded. Output files rank events by seed match length and observed off-target log fold-change.

### Candidate off-target nomination by transcriptional downregulation

For each guide, knockdown of the intended target gene was assessed by computing log2 fold-change on pseudobulk values relative to non-targeting guides, with a pseudocount of 0.01 added to both the numerator and denominator. For each neighboring guide, the expression of that neighbor’s target gene was checked to determine whether the current guide downregulates the neighbor guide target gene.

### Reanalysis of K562 GWPS data

For figure 1f, the GWPS dataset is loaded in Scanpy in disk-backed mode^49^. All cells receiving guides targeting either *SMG5*, *SMG7*, *UPF1*, *UPF2*, or *CLOCK* are loaded into memory, as well as a random sampling of 500 non-targeting control cells. A Wilcoxon rank sum test was performed between the ‘CLOCK’ and ‘non-targeting’ cells using the ‘rank_genes_groups’ function in Scanpy. The top 12 genes with greater than 1 log2 fold-change and a p-value of less than 0.01 were plotted after normalizing and scaling raw counts. Figure 1b displays clustered correlation of principal components of pseudobulk gene expression values with the pseudobulked non-targeting control values subtracted. Pairwise Pearson correlation coefficients were computed across the low-dimensional embedding to generate a guide-by-guide similarity matrix.

### Off-target workflow of GWPS datasets

Each GWPS dataset was downloaded from the links listed under ‘Data Availability.’ For the VIPerturb-seq dataset, the downloaded Seurat object was converted to an AnnData object. The VIPerturb Seurat object was loaded using Seurat v5.4 and R version 4.5^50^. Then, raw counts were extracted from the ‘counts’ layer of the ‘RNA’ assay using the ‘GetAssayData’ function. Raw counts were written in market matrix format using the ‘Matrix::writeMM’ function. The metadata slot of the Seurat object was additionally saved as a CSV. The saved matrix was loaded in python using the ‘mmread’ function from SciPy^51^. All other objects were used exactly as they were downloaded from the provided sources.

We used the same parameters for each of the four datasets analyzed. In each case, GENCODE v47 TSS annotations were used along with the hg38 reference genome for seed matching. We specified a minimum seed match cutoff of 5 bp. For pseudobulk guide dimensionality reduction and neighborhood finding, the top 5,000 features were considered following normalization. PCA was run with 100 principal components. Using this embedding, 20 nearest neighbors were identified by Euclidean distance and extended to include asymmetrical neighbors. To reduce noise in guide-level pseudobulk profiles, guides with less than 20 assigned cells were filtered from all analyses.

### Motif enrichment plots

Bar plots displaying the 2 bp motif enrichment in the 10 bp seeds of candidate off-target guides with ≥10 bp seed matches and a negative off-target log2 fold-change across all four GWPS datasets (Replogle, Xaira, Zhu & Dann, and Bradu & Blair). Enrichment was calculated as the ratio of observed to expected motif frequency, where observed frequency was derived from the 10 bp seeds of candidate off-target guides and expected frequency was derived from the equivalent seed regions across the entire genome-wide guide library. Sequence logo plots were computed using the same candidate off-target guides. The frequency of each base pair at each position was computed for the candidates as well as all guides in the library. The base frequencies for the whole library were then subtracted from the candidate off-target frequencies and plotted using the logomaker package.^52^

### TSS enrichment plots

To assess the proximity of candidate off-target editing events to transcription start sites (TSS), the genomic position of each candidate off-target site was recorded. The distance from each off-target site to its nearest TSS was calculated by intersecting off-target coordinates with TSS annotations derived from GENCODE v47^53^. Distances were computed relative to the TSS, accounting for transcript strand orientation such that negative values indicate positions upstream of the TSS and positive values indicate positions downstream.

### Recorded output

The workflow generates a single summary CSV file as output. Each row of the CSV documents an event between a guide and off-target gene. The log2 fold-change of both the on-target and off-target genes is recorded, relative to non-targeting control guides. If a seed match of 5 or more bp is detected in the 4,000 bp window surrounding the off-target TSS, the location and length of the match is recorded. If no value is listed, then that event only contains transcriptional evidence of off-target repression. Output CSVs are sorted first by the length of the seed match and secondarily by the log2 fold-change of the candidate off-target gene, such that higher confidence off-targets are found at the beginning of the output file.

### Repression significance analysis

To ensure significance of our pseudobulked log2 fold-change measures, we ran single cell two-sided Wilcoxon rank-sum tests to evaluate whether candidate off-target genes with seed length matches of a given length were statistically significant. Tests were run using the ‘mannwhitneyu’ function from SciPy stats^51^. To maintain efficiency, 500 candidate events for seed match lengths between 5 and 9 were randomly sampled, while all events of 10 or more bp were kept. Candidate off-target events were labeled as significant if the log2 fold-change was negative and the p-value was less than 0.01.

### Reanalysis of Schmidt et al, Belk et al, and Datlinger et al data

For Schmidt et al, supplementary table 1 was downloaded^39^ and the sheets containing MaGECK^54^ output for each of the four plotted screens were extracted. For CRISPRa screens, we plotted the -log10 value of ‘pos|score’ and plot ‘pos|rank’ on the x-axis. For CRISPRi screens, we plotted the -log10 value of ‘neg|score’ and plot ‘neg|rank’ on the x-axis. For the *CALCOCO2* and *LAT* guide-level quantification, we downloaded fastqs for GEO samples^55^ with accession numbers: GSM5290228, GSM5290229, GSM5290240, and GSM5290241 using fasterq-dump from the SRA toolkit. Guide counts for each of the *CALCOCO2* and *LAT* Dolcetto Set A^12^ guides were determined by grep search and normalized based on the total read number in the fastq. Belk et al data was downloaded from supplementary table 1 and the rank ordered ‘casTLE_score’ column was plotted^37^. For the Datlinger et al data, we plotted the -log10 value of the ‘neg|score’ along the y-axis and ‘neg|rank’ on the x-axis^38^. In each of these plots, we highlight *CD3E, CD3G, CD247, LAT, CD3D, LCP2, ZAP70, PLCG1*, and *VAV1* as canonical regulators of the TCR pathway and *LRBA, APPL2, WDR53,* and *CALCOCO2* as recently nominated putative regulators.

## CODE & RESOURCE AVAILABILITY

Seed matching web application:

https://crispr-seed-finder.vercel.app/ GitHub repository for pipeline:

https://github.com/AustinHartman/perturb_seed GitHub repository to reproduce analysis:

https://github.com/AustinHartman/perturb_seed_reproduce

## DATA AVAILABILITY

Each GWPS dataset is made available by the authors at the following locations: Replogle K562 GWPS: https://doi.org/10.25452/figshare.plus.20029387

Xaira HCT116 GWPS: https://doi.org/10.25452/figshare.plus.29190726 Zhu & Dann CD4+ T GWPS:

https://virtualcellmodels.cziscience.com/dataset/genome-scale-tcell-perturb-seq Bradu & Blair genome-wide VIPerturb-seq: https://zenodo.org/records/18460279

## ACKNOWLEDGEMENTS

We thank members of the Roth and Satpathy labs for useful discussion. A.T.S. was supported by the Cancer Research Institute, the Parker Institute for Cancer Immunotherapy (PICI), and the Mark Foundation. The Marson laboratory has received research support from the PICI, the Emerson Collective, Arc Institute, Juno Therapeutics, Epinomics, Sanofi, GlaxoSmithKline, Gilead and Anthem and reagents from 10x, Ultima, Genscript, Illumina and Cellanome.

## AUTHOR CONTRIBUTIONS

A.H. conceptualized the study. A.H. wrote and edited the manuscript with input from all authors.

A.H. performed analysis. M.B. built the web application with assistance from J.D.B. and A.H. J.D.B., R.S., A.T.S., and T.L.R. guided the analysis.

---

**Extended Data Figure 1:**
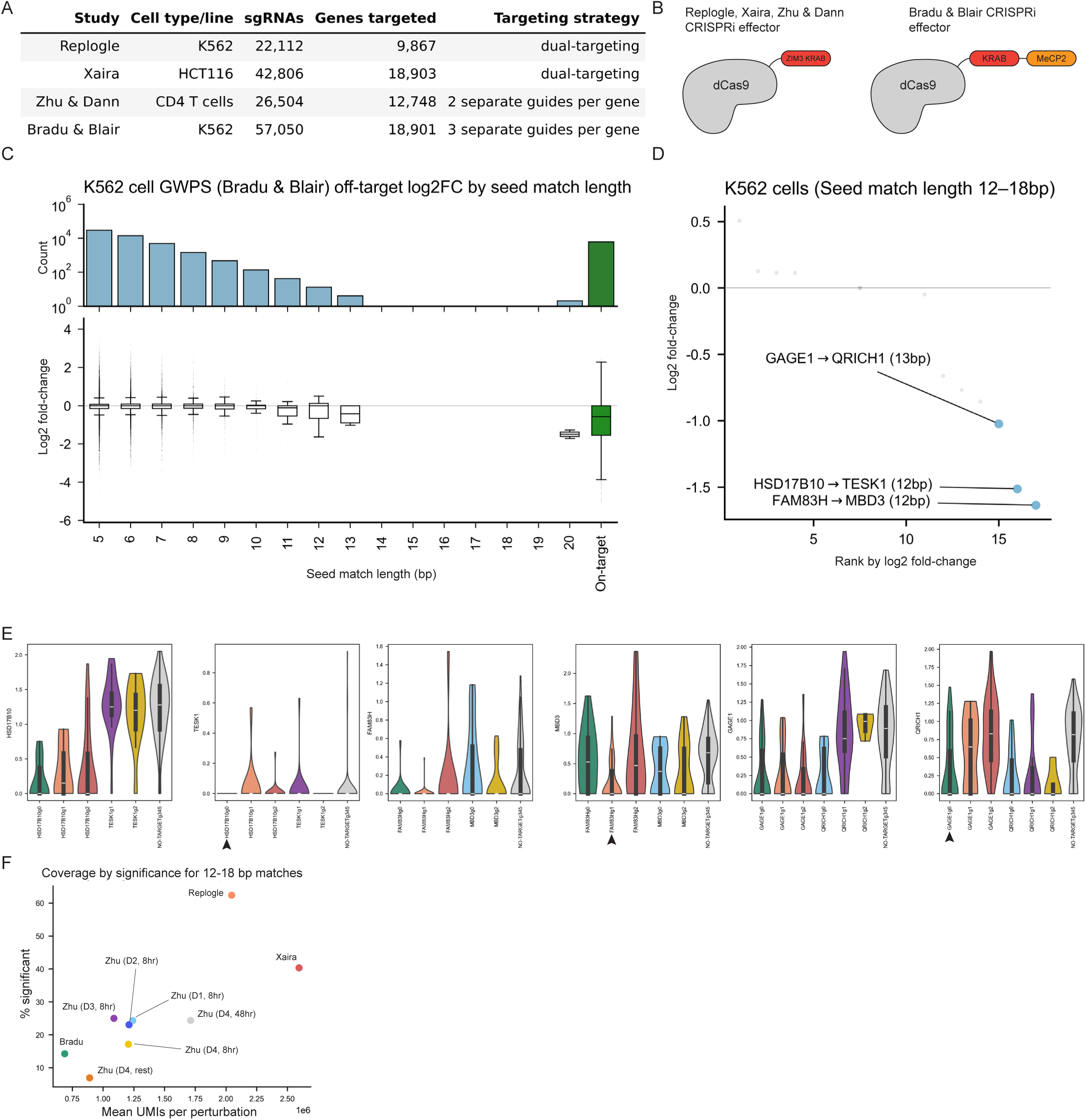
GWPS dataset overview and application to VIPerturb-seq dataset. **a.** Overview of the guide libraries and cells used in each study. **b.** Graphic of the CRISPRi architecture used across four separate GWPS datasets. **c.** Measured log2 fold-change for candidate off-target guides at the predicted off-target guide split by the length of seed sequence match in a GWPS dataset in K562 cells (Bradu & Blair). Seed matches of 19 or 20 bp indicate off-target repression of a neighboring gene, while the others represent possible off-target events. The final column of the plots display measured log2 fold-change for guides at their intended targets. **d.** Log2 fold-change rank plot showing off-target 12-18 bp seed matches. The number of off-target events identified in a dataset should not be interpreted as a measure of data quality, as it reflects many factors such as the number of guides, guide architecture, number of cells sequenced, number of cells per guide, and active pathways within cells. The three top candidate off-target guides based on log2 fold-change were colored in blue. **e.** Violin plots of three top off-target candidates. Up to 3 independent guides target each gene. The guides annotated with arrows indicate the guides with predicted off-target activity of the gene plotted. **f.** Scatter plot displaying the fraction of significant off-target seed matches of 12-18 bp using a Wilcoxon rank sum test by the mean number of UMIs per perturbation.

**Extended Data Figure 2:**
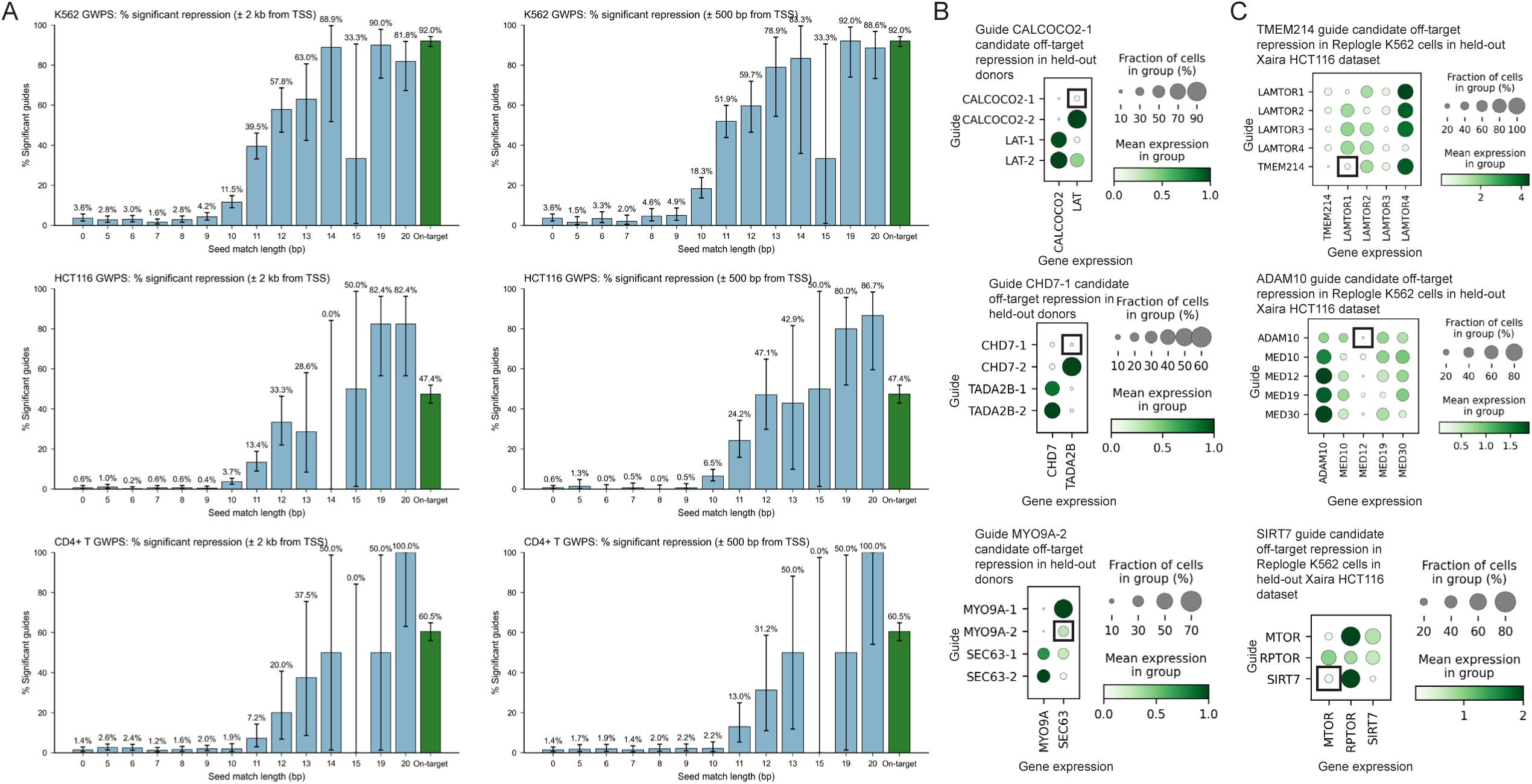
Validation of CRISPRi off-target repression across independent donors and cell lines. **a.** Plot displaying the fraction of significant off-target seed matches using a Wilcoxon rank sum test by seed match length. To reduce the number of tests run, 500 events were sampled for seed match lengths 0 through 9 as well as the ‘On-target’ category colored in green. Clopper-Pearson confidence intervals were displayed for each bar with a 0.95 confidence level. **b.** Dot plots displaying candidate off-target repression identified in CD4+ T cells donor 1 dataset in three additional held-out donors (donor 2, 3, and 4) each stimulated for 8 hours. Each gene was targeted by two independent guides suffixed with “-1” or “-2” and the predicted off-target guide and gene combination is highlighted with a black box. **c.** Dot plots displaying candidate off-target repression events identified from GWPS data in K562 cells in a separate GWPS experiment using the same guide pairs in HCT116 cells, a colorectal carcinoma cell line. The predicted off-target guide and gene combination is highlighted with a black box.

**Extended Data Figure 3:**
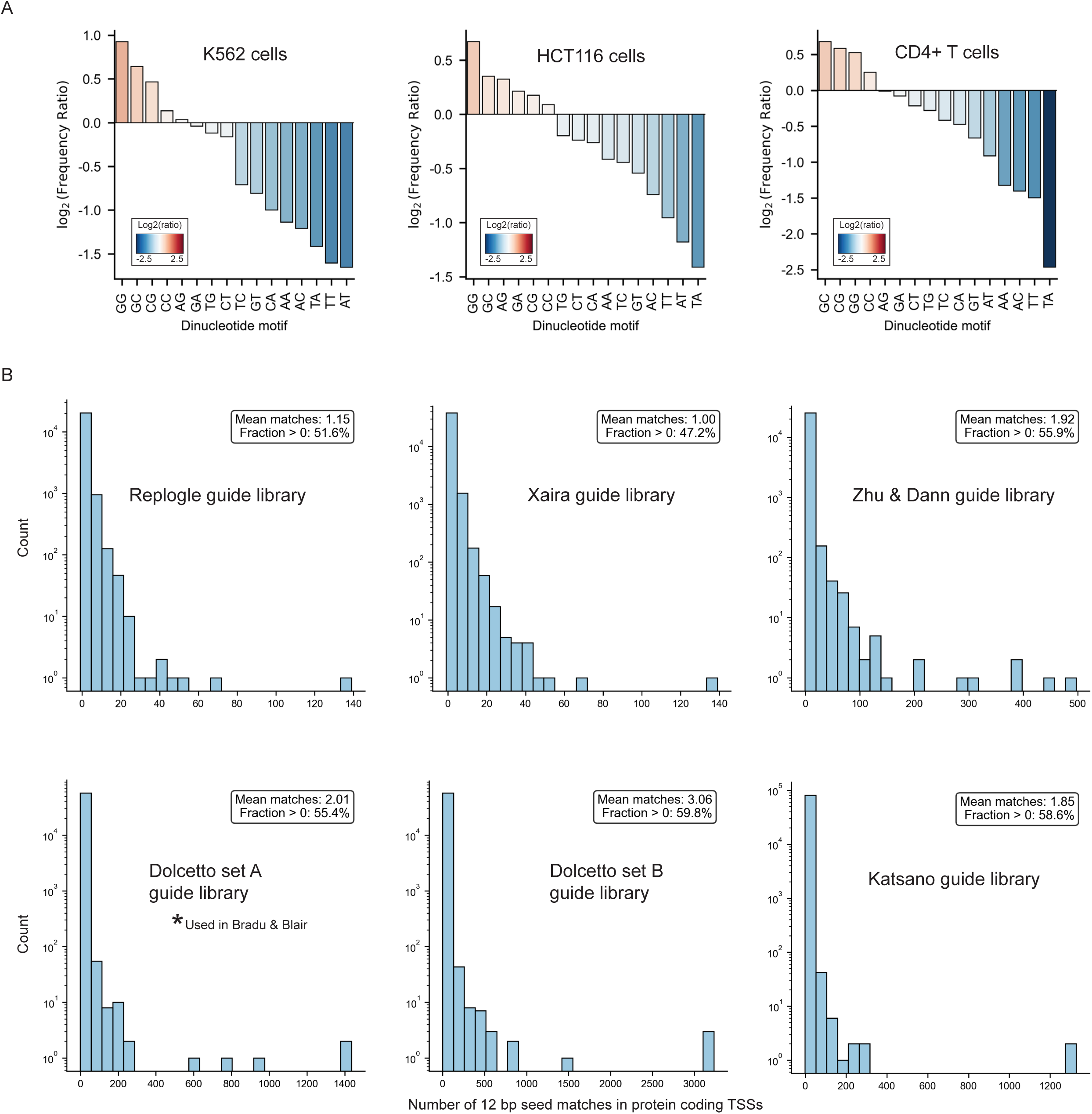
Sequence features and prevalence of off-target seed matches in genome-wide CRISPRi guide libraries. **a.** Bar plots displaying log2 dinucleotide enrichment in the 10 bp seeds of candidate off-target guides with ≥10 bp seed matches and negative log2 fold-change of the candidate off-target gene across three GWPS datasets. Enrichment values were determined by computing the frequency of each motif in all candidate off-target seeds versus the seeds in the whole guide library. Values greater than zero indicate enrichment in the candidate off-target guides. **b.** Off-target 12 bp TSS seed matches across various genome-wide CRISPRi guide libraries. A value of 1 is subtracted from each guide count, given the assumption that one count per guide represents the alignment to the on-target locus. Non-targeting control guides were filtered and only protein coding TSSs were considered. Protein coding MANE transcript TSSs annotations were used.

**Extended Data Figure 4:**
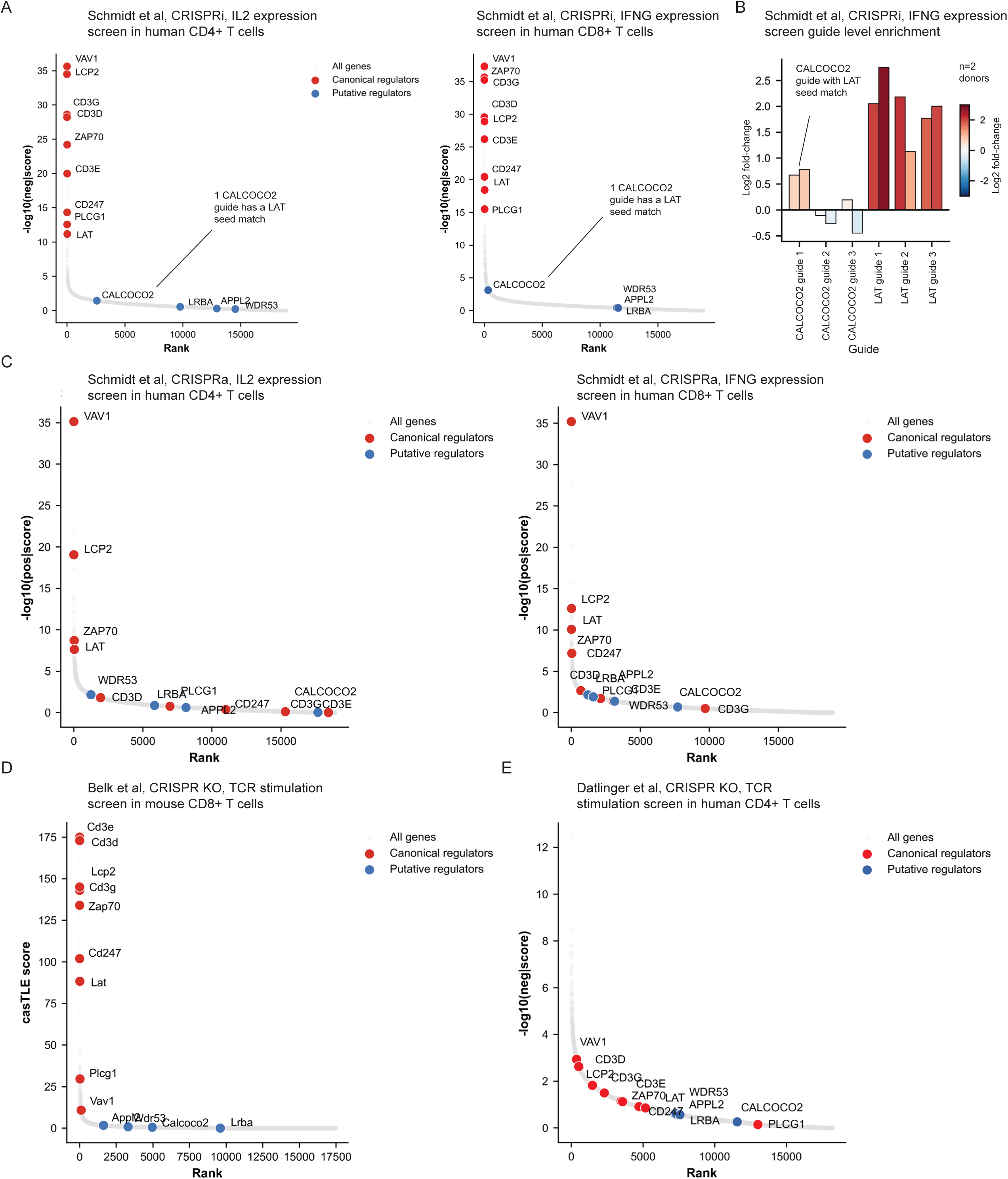
Specific enrichment of canonical TCR regulators across genome-wide CRISPR screens in primary human and mouse T cells. **a.** Rank plots of MAGeCK negative enrichment scores from CRISPRi experiments for IL2 expression in CD4+ T cells and IFNG expression in CD8+ T cells where each gene is targeted by 3 to 6 independent guides from the dolcetto library. **b.** Log2 fold-change enrichment for IFNG-low relative to IFNG-high CD8+ T cells for six guides from Dolcetto Set A targeting *CALCOCO2* and *LAT*. **c.** Rank plots of MAGeCK negative enrichment scores from CRISPRa experiments for IL2 expression in CD4+ T cells and IFNG expression in CD8+ T cells where each gene is targeted by 3 to 6 independent guides from the Calabrese library. **d.** CRISPR-Cas9 knockout screen in CD8+ mouse T cells following 8 days of anti-CD3 stimulation. **e.** CRISPR knockout screen in human CD4+ T cells following anti-CD3/CD28 stimulation. Rank plots display negative enrichment scores computed using MaGECK.

